# Purification of Human β- and γ-actin from Budding Yeast

**DOI:** 10.1101/2022.08.17.504301

**Authors:** Brian K. Haarer, Morgan L. Pimm, Ebbing P. de Jong, David C. Amberg, Jessica L. Henty-Ridilla

## Abstract

Biochemical studies of human actin and its binding partners rely heavily on abundant and easily purified α-actin from skeletal muscle. Therefore, muscle actin has been used to evaluate and determine the activities of most actin regulatory proteins and there is an underlying concern that these proteins perform differently with actin present in non-muscle cells. To provide easily accessible and relatively abundant sources of human β- or γ-actin (i.e., cytoplasmic actins), we developed *Saccharomyces cerevisiae* strains that express each as their sole source of actin. Both β- or γ-actin purified in this system polymerize and interact with various binding partners, including profilin, mDia1 (formin), fascin, and thymosin-β4 (Tβ4). Notably, Tβ4 and profilin bind to β- or γ-actin with higher affinity than to α-actin, emphasizing the value of testing actin ligands with specific actin isoforms. These reagents will make specific isoforms of actin more accessible for future studies of actin regulation.

## Introduction

The actin cytoskeleton is an essential, highly conserved, and abundant component of cells. Simple eukaryotes tend to express a single actin isoform, while humans display tissue and cell specific expression patterns. Although closely related, actin isoforms can subtly differ in biochemical properties related to polymer formation, nucleotide hydrolysis and exchange, and interactions with one or more essential regulatory proteins (Allen et al., 1996; Moradi et al., 2017; Namba et al., 1992; Perrin and Ervasti, 2010). Humans express six actin isoforms: α1, α2, α-cardiac, and γ2, which occur predominantly in skeletal, cardiac, and smooth muscle cells, and β and γ1 (γ henceforth) found predominantly in non-muscle cells and considered cytoplasmic isoforms of actin. Human β- and γ-actin are structurally divergent (Arora et al., 2023), yet differ by only four amino acids located within their first ten amino acids (Figure 1A). Various muscle tissues are the most common sources of actin for use in biochemical studies. Actin purified in this manner is present as a mixture of muscle isoforms and minor amounts of β or γ-actin. Characterizing specific isoforms of actin has been limited by accessibility and is critically important for understanding mechanisms of disease (Parker et al., 2020). Thus, it has been challenging to discern the biochemical properties of either β- and γ-actin in biochemical assays.

**Figure 1.**
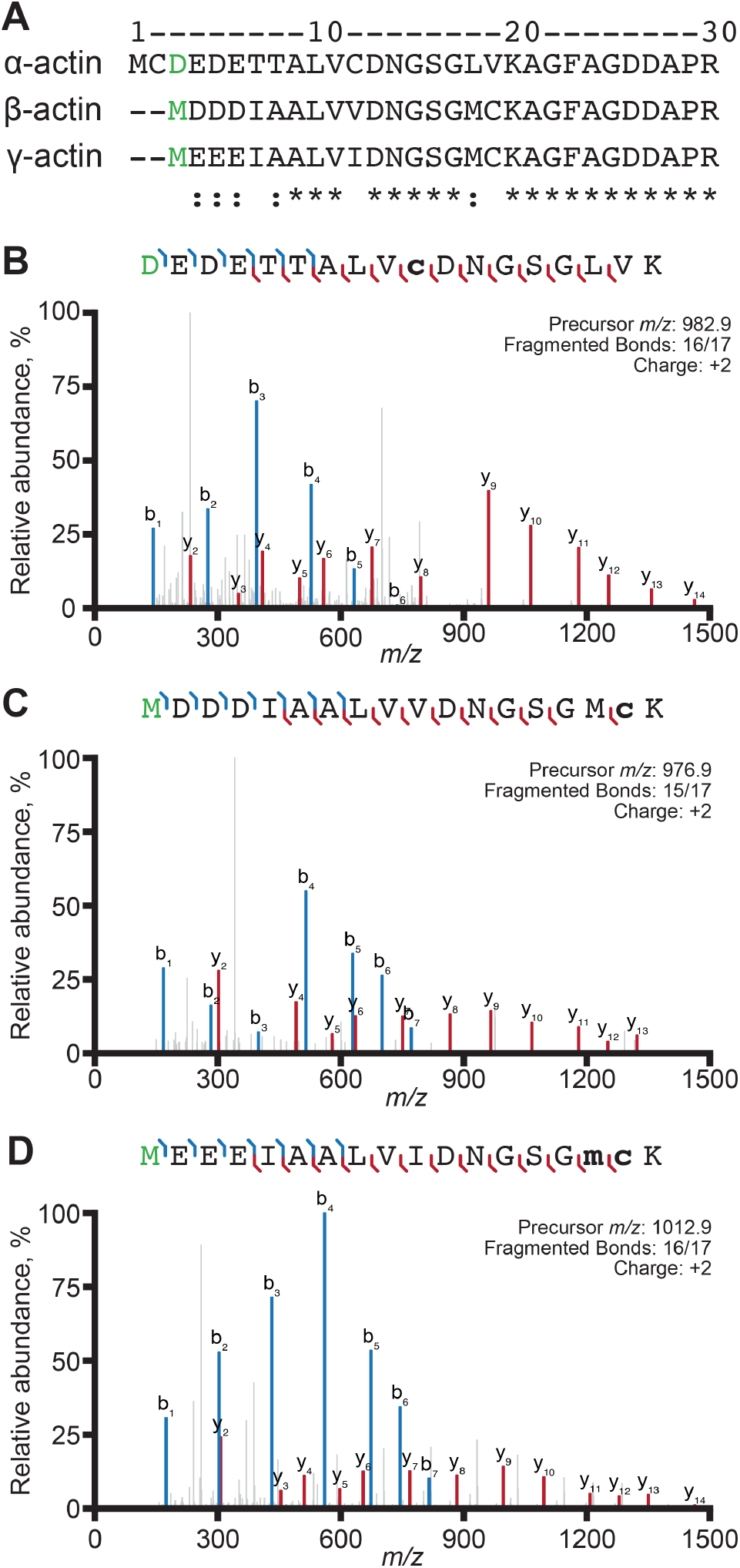
β-actin and γ-actin are acetylated on the N-terminal methionine. **(A)** Sequence alignment of human α1-actin, β-actin, and γ1-actin. Spectra of acetylated (green) N-terminal peptides as indicated for **(B)** α-actin (RMA), **(C)** β-actin, or **(D)** γ-actin. Bold lower-case letters: m, oxidized methionine; c, carbamidomethyl cysteine.

Producing recombinant human actin outside of eukaryotic cells is difficult due to the complex network of chaperones needed to properly fold actin, however several systems have been developed (Geissler et al., 1998; Grantham, 2020; Millán-Zambrano and Chávez, 2014; Schafer et al., 1998; Valpuesta et al., 2002). The pCold system permits the bacterial synthesis of recombinant tagged β-actin (Tamura et al., 2011). Actin isoforms expressed and purified from popular eukaryotic systems produce relatively large quantities of biochemically active actin (Bergeron et al., 2010; Bookwalter and Trybus, 2006; Ohki et al., 2009; Rutkevich et al., 2006; Yamashiro et al., 2014). While these and related purification methods have been adopted for various studies of normal and mutant versions of actin, preparations are often contaminated with low amounts (5-15%) of host actin (Hundt et al., 2014; Müller et al., 2012; Müller et al., 2013; von der Ecken et al., 2016). Other systems use a combination of affinity tags and a direct fusion to the actin monomer binding protein thymosinβ4 (Tβ4) to prevent the spurious polymerization or aggregation of recombinant actin and to facilitate isoform specific purification (A et al., 2020; Hatano et al., 2018; Hatano et al., 2020; Kijima et al., 2016; Lu et al., 2015; Noguchi et al., 2007). With the addition of NAA80 or SETD3 modifications this approach permits the isolation of actin in specific post translationally modified states, including N-acetylation, N-arginylation, and methylation of a conserved histidine residue (Arora et al., 2023; Hatano et al., 2018; Hatano et al., 2020). Additional approaches even permit the close preservation of native secondary modifications (Ceron et al., 2022). While each system requires post-purification processing, they provide a source of non-muscle, normal or mutant (including non-polymerizable), actin isoforms.

We have taken a different approach to generate pure human β- or γ-actin, engineering the yeast *Saccharomyces cerevisiae* to produce either isoform as their only source of actin. This technique was first pioneered to purify chicken β-actin from yeast, relying on hydroxylapatite chromatography to separate host and recombinant actin (Karlsson, 1988). A follow-up study showed that yeast could survive, albeit not well, with this as their sole actin source, although β-actin was not purified from these yeast strains (Karlsson et al., 1991). We have taken codon-optimized genes for human β- and γ-actin and expressed them in yeast lacking the resident actin gene, *ACT1*. The resulting strains grow considerably slower than wild type yeast yet provide significant, low-cost, and easily obtainable yields of homogeneous β- or γ-actin. While actin purified by this approach does not undergo standard mammalian N-terminal processing, each actin isoform polymerizes and interacts with several actin-binding proteins including profilin, Tβ4, the formin mDia1, and fascin. Thus, these engineered strains provide convenient sources of pure β- or γ-actin for biochemical studies where conventional α-actin sources may be unsuitable.

## Methods

### Reagents and supplies

Unless otherwise specified, chemicals and supplies were purchased from Fisher Scientific (Pittsburgh, PA). DNaseI for affinity columns was purchased from Worthington Biochemical (Lakewood, NJ). Cloning reagents were obtained as follows: Restriction enzymes and DNA ligase (New England Biolabs; Ipswich, MA); Primestar HS DNA polymerase (Takara Bio USA; San Jose, CA); Oligonucleotides (Eurofins Genomics; Louisville, KY).

### Plasmid and strain construction

Human α1-actin (*ACTA1*; NCBI Gene ID: 58) and β-actin (*ACTB*; NCBI Gene ID: 60) (Figure S1) genes flanked by BamHI and HindIII sites were codon-optimized for expression in budding yeast (Figures S1A and S1B) (GenScript; Piscataway, NJ). Codon-optimized γ1-actin (*ACTG1*; NCBI Gene ID: 71) was generated from the *ACTB* sequence with the following mutagenic primer (5 ‘- CTCGGATCCATGGAAGAAGAAATTGCTGCATTGGTTATTGATAATGGTTCTGGCATGTG-3 ‘). The TDH3 promoter was PCR amplified as an EcoRI-BamHI fragment from yeast genomic DNA with 5 ‘- GTGAGAATTCTCAGTTCGAGTTTATCATTA-3 ‘ and 5 ‘-CTCGGATCCTTTGTTTGTTTATGTGTGTTT-3 ‘. TDH3 and actin genes were inserted into YEp351. The yeast strain expressing only β-actin (BHY845) is a haploid segregant of an *ACT1/act1*Δ heterozygous diploid that carried the β-actin expression plasmid. The yeast strain that expresses only γ1-actin (BHY848) was generated by introducing the γ1-actin expression plasmid into a haploid *act1*Δ strain carrying the yeast *ACT1* gene on a URA3-based plasmid. We used 5-fluoroorotic acid (FOA) to select for yeast that lost the *ACT1* (*URA3*) plasmid but remained viable. We were unable to recover yeast expressing human α1-actin, consistent with previous reports (McKane et al., 2005; McKane et al., 2006). We re-isolated and sequenced plasmids from β- and γ-actin strains at the time of harvest to confirm sequence fidelity. Each actin gene sequence was identical to the starting yeast-optimized *ACTB* and *ACTG1* genes.

### Protein purification

Human β- and γ-actin were prepared as described in Aggeli et al. (2014), with the following modifications: yeast were grown in 1 L of YPD (1% yeast extract, 2% bactopeptone, 2-4% glucose), harvested by centrifugation at 8000 × *g* for 7 min, washed in 25 mL 10 mM Tris (pH 7.5), 0.2 mM CaCl_2_. Pellets were collected by centrifugation at 4000 × *g* for 10 min and stored frozen at -80 °C. Pellets were suspended in 20 mL G-buffer (10 mM Tris (pH 7.5), 0.2 mM CaCl_2_, 0.5 mM ATP, 0.2 mM DTT) supplemented with protease inhibitors (0.1 mM PMSF and 1:500 Cocktail IV (Calbiochem; San Diego, CA), and lysed by two passes through a French press at 1000 psi. Lysates were bound to 3 mL DNaseI-Sepharose, then beads were washed in five column volumes of each of the following: (1) 10% formamide in G-buffer, (2) 0.2 M NH_4_Cl in G-buffer, and (3) G-buffer alone. Actin was eluted with 50% formamide in G-buffer, and then immediately diluted with 1-2 mL G-buffer and dialyzed overnight against 2 L G-buffer. To remove contaminating proteins, actin underwent at least two polymerization-depolymerization cycles alternating between 0.6 M KCl and pipetting with dialysis against G-buffer.

Rabbit muscle actin (RMA; α-actin), mDia1(FH1-C) (amino acids 571-1255), GFP-thymosin-β4, profilin-1, and fascin used in TIRF or anisotropy assays were purified as described (Jansen et al., 2011; Liu et al., 2022; Pimm et al., 2022). Actin was labeled with Oregon Green iodoacetamide or Alexa Fluor NHS-Ester (Aggeli et al., 2014; Hertzog and Carlier, 2005; Kuhn and Pollard, 2005). Proteins were aliquoted, flash-frozen in liquid nitrogen, and stored at -80 °C. Proteins used in TIRF assays are shown in Figure S6.

### Mass spectrometry (MS) analysis

Sample preparation, operation, and MS analyses were performed by the Upstate Medical University Proteomics and Mass Spectrometry Core Facility. Samples (15-30 µg) were treated with 5 mM TCEP (tris(2-carboxyethyl)phosphine) and then 15 mM iodoacetamide for 30 min in the dark. Trypsin digestion was performed at 37 °C with 0.66 µg Trypsin Platinum (Promega, Madison, WI) overnight. Samples were acidified and then desalted using 2-core MCX stage tips (Rappsilber et al., 2003). Peptides were eluted with 75 µL of 65% acetonitrile (ACN) with 5% ammonium hydroxide, and dried. Samples were dissolved in 65 µL 2% ACN and 0.5% formic acid in water, then 0.5 µg was injected onto a pulled tip nano-LC column held at 50 °C, with 100 µm inner diameter packed to 32 cm with 1.9 µm, 100Å, core shell Magic2 C18AQ particles (Premier LCMS, Penn Valley, CA). Peptides were separated with a 3 – 28% ACN gradient over 60 min, followed by an increase to 85% ACN over 5 min. The inline Orbitrap Lumos was operated at 2.3 kV in data-dependent mode with a cycle time of 2.5 s. MS1 scans were collected at 120,000 resolution with a maximum injection time of 50 ms. Dynamic exclusion was applied for 15 s. Stepped HCD fragmentation of 34, 38, and 42% collision energy was used followed by two MS2 microscans in the Orbitrap at 15,000 resolution with dynamic maximum injection time.

The core facility used SequestHT in Proteome Discoverer (version 2.4; Thermo Scientific) to search three MS databases: *S. cerevisiae* (Uniprot, 6816 entries, retrieved 2017), Rabbit *O. cuniculus* (Uniprot, 41462 entries, retrieved 2023) a list of common contaminants (Thermo Scientific, 298 entries), and a custom list of human β-actin, γ-actin, and rabbit α-actin. Additional entries for each actin were made with truncations of the first one, two and three N-terminal residues. Enzyme specificity was set to semi-tryptic with up to 2 missed cleavages. Precursor and product ion mass tolerances were set to 10 ppm and 0.02 Da. Cysteine carbamidomethylation was set as a fixed modification and the following modifications were set as variable: Methionine oxidation, protein N-terminal acetylation, peptide N-terminal arginylation, lysine acetylation or methylation of lysine, histidine, and arginine. The output was filtered using the Percolator algorithm. We visualized datasets with: http://www.interactivepeptidespectralannotator.com/PeptideAnnotator.html (Brademan et al., 2019). Full data sets are deposited at the PRIDE Database (ProteomeXchange identity: PXD040174).

### F-actin sedimentation assays

Polymerization of 1 or 2 µM actin was induced with the addition of concentrated (20×) F-buffer (final 1× concentration: 10 mM Tris (pH 7.5), 25 mM KCl, 4 mM MgCl_2_, 1 mM EGTA, 0.5 mM ATP). Samples were incubated for 30 min at room temperature, then subjected to centrifugation at 200,000 × *g* for 30 min at 20 °C. Supernatants were removed and pellets and supernatant samples were brought to equal volumes in protein loading buffer. Equal volumes were loaded on SDS-polyacrylamide gels and stained with Coomassie. Indicated concentrations of profilin-1 were mixed with 2 µM actin monomers in G-buffer and then incubated for 30 min at room temperature. Actin polymerization was then induced with F-buffer for an additional 30 min prior to centrifugation as above.

### Fluorescence-based actin polymerization assays

Actin isoform co-polymerization reactions were assessed by epifluorescence microscopy (Zeiss Imager.Z1, Oberkochen, Germany). 2 µM unlabeled or OG-labeled β- or γ-actin were polymerized in F-buffer, either individually or 1:1 mixture, then bound to 1.1 µM rhodamine-phalloidin. Samples were viewed individually or as mixtures bound to phalloidin prior to mixing. Samples were flowed into homemade slide chambers prepared by laying a cover glass onto a slide with two intervening strips of double-sided tape, then visualized with DsRed and FITC filter sets. Views of were obtained near fluid/air boundaries.

### Total Internal Reflection Fluorescence (TIRF) microscopy assays

We prepared and visualized TIRF imaging chambers on a DMi8 TIRF microscope (Leica Microsystems; Wetzlar, Germany) as in Henty-Ridilla (2022), with the following modifications: reactions were executed in a different TIRF buffer (20 mM imidazole (pH 7.4) 50 mM KCl, 1 mM MgCl_2_, 1 mM EGTA, 0.2 mM ATP, 10 mM DTT, 40 mM glucose, and 0.25% methylcellulose (4000 cP)), with minimal (5-7%) labeled-RMA. Some TIRF experiments with β- or γ- actin were supplemented with 10% RMA to visualize filaments, as noted in figure legends. Experiments performed with fascin used unlabeled actin isoforms and were visualized with 130 nM Alexa-488 phalloidin. Filament elongation rates (subunits s^-1^µM^-1^) were determined by measuring filament lengths from at least five frames, with conversion factor of 370 subunits/µm (Kuhn and Pollard, 2005). Total filament or bundle length was calculated using the FIJI Ridge Detection plugin with settings that minimized background signals but permitted the detection of faint filaments without image saturation and applied identically to all images (Steger, 1998; Wagner et al., 2017). Skewness parameter was measured from FIJI measurements (Higaki et al., 2010; Khurana et al., 2010; Schindelin et al., 2012).

### Fluorescence polarization assays

Fluorescence polarization determination of GFP-thymosin- β4 (Tβ4) binding to actin (Liu et al., 2022) was carried out in 1× PBS (pH 8.0) supplemented with 150 mM NaCl. 10 nM β- or γ-actin were mixed with concentrations (0.1 pM to 10 µM) of GFP-Tβ4 and incubated at room temperature for 15 min. Fluorescence polarization was measured in a plate reader equipped with monochromator, with excitation at 440 nm and emission intensity detection at 510 nm (bandwidth set to 20 nm) (Tecan; Mannedorf, Switzerland). Three technical replicates were carried out on the same plate.

### Data analysis, statistics, and availability

GraphPad Prism (version 9.5.0; GraphPad Software, San Diego, CA) was used to plot all data and perform statistical tests. All experiments were repeated at least 3 times. Individual data points are presented as dots in each figure and histograms represent means ± SE (unless noted otherwise). All one-way ANOVA tests performed compared all means with Tukey post-hoc analysis and passed tests for normality. P-values presented are those from ANOVA tests across tested conditions (under the line), except for Figure S5 where all comparisons are listed. The threshold (p ≤ 0.05) was used to determine significance throughout this work. Specific comparisons are described in each figure legend. Figures were made in Adobe Illustrator 2023 (version 27.1.1; Adobe, San Jose,CA). MS datasets are deposited at the PRIDE Database (ProteomeXchange identity: PXD040174). All other datasets used to generate each figure are available here: https://doi.org/10.5281/zenodo.7641497.

## Results

We expressed codon-optimized human cytoplasmic β- or γ- actin in *Saccharomyces cerevisiae* strains lacking the sole yeast actin gene (*ACT1*). The strain expressing human β-actin grew slowly (Figure S2A) and displayed heterogeneous morphology (Figure S2B-D), similar to yeast expressing chicken β-actin (Karlsson et al., 1991). The strain expressing γ-actin grew even more slowly (Figure S2A). However, either strain could be maintained without additional selection for the essential plasmid-borne actin genes. To confirm the identity and assess the posttranslational state of either isoform, each recombinant actin was purified by affinity chromatography and subjected to mass spectrometry. This analysis showed 89% (β-actin), 95% (γ-actin), and 98% (α-actin; RMA) coverage of the predicted peptide profiles. We took special note of the state of N-terminal peptides, which are the only peptides expected to differ between β- and γ-actin (Figure 1A). We detected N-terminally acetylated β- and γ-actin (at Met1)(Figure 1C and 1D), although not all intact peptides were N-acetylated. We also detected non-acetylated β- and γ-actin peptides truncated by 1 to 5 amino acids (Table S1, Table S2, and Data File S1) For comparison, all N-terminal peptides on α-actin (control) were truncated by 2 to 6 amino acids, present with and without acetylation on Asp3 (Figure 1B, Table S1, Table S2, and Data File S1).

Two other important modifications of metazoan versions of actin include the methylation of His73 which may affect Pi release following ATP hydrolysis (Terman and Kashina, 2013), and Lys84, which may block myosin binding (Li et al., 2013). Each of these modifications were present in α- actin (control) dataset but were absent in the β- and γ-actin spectra (Table S1). Finally, N-terminal arginylation of β- actin on Asp3 can influence cellular and biochemical functions (Chin et al., 2022; Karakozova et al., 2006). However, N-arginylation of β- and γ-actin on the 3rd residue cannot be reliably set apart from N-acetylation on the 2nd residue (Tables S1 and S2)(Karakozova et al., 2016; Drazic et al., 2022). Not surprisingly, the modification profiles of β- and γ-actin reflect processing that normally occurs for yeast actin. Yeast processing includes the acetylation of N-terminal Met1 without further processing and also lacks additional methylating enzymes (Figure 1; Tables S1 and S2)(Kalhor et al., 1999).

The ability of either β- or γ-actin to support yeast growth suggests protein functionality (Figure S2). To explore this idea further, we used classic biochemical assays to characterize each cytoplasmic actin. We began by assessing the ability of the β- and γ-actin to polymerize into filaments with conventional pelleting assays (Figure 2A-B). Polymerization of each actin was indistinguishable, with filaments pelleting to similar degrees at multiple concentrations (Figure 2A-C). Each actin remained in the supernatant under controls performed in G-buffer (Figure S3). To assess whether β- and γ-actin could co-polymerize, we performed epifluorescence microscopy to visualize actin filaments polymerized before or after mixing unlabeled and Oregon Green (OG)-labeled β- or γ-actin bound to rhodamine(Rh)-phalloidin (Figure 2D-E). Rh-phalloidin stabilization of actins polymerized before mixing resulted in a mixed population of filaments including Rh-phalloidin-decorated unlabeled actin (only visible in the Rh-phalloidin channel) and OG-labeled filaments co-labeled with Rh-phalloidin (visible in both channels) (Figure 2D-E, top). In contrast, mixing of actins prior to polymerization, followed by Rh-phalloidin stabilization, produced uniformly labeled actin filaments in both wavelengths, consistent with co-polymerization (Figure 2D-E, bottom). Notably, filament co-labeling occurred regardless of which actin is OG-labeled, consistent with co-polymerization of β- and γ-actin. To further assess the actin assembly properties of β- and γ-actin, we used total internal reflection fluorescence (TIRF) microscopy to directly observe single actin filament polymerization (Figure 3A)(Movie 1). Unlabeled 1 µM β-actin, γ-actin, or control α-actin (RMA) was visualized with 10% fluorescently labeled RMA (Figure 3A)(Chin et al., 2022; Hatano et al., 2018). Under these conditions, α-, β-, or γ-actin show similar means of nucleation ranging from 25.5 ± 5.5 to 59.7 ±19.3 filaments per field of view (p ≥ 0.45)(Figure 3B). Control α-actin filaments (RMA) elongated at a rate of 10.0 subunits s^-1^ µM^-1^ ± 0.3 (SE), consistent with other studies (Figure 3C)(Liu et al., 2022; Pimm et al., 2022). β-actin and γ- actin filaments polymerized at mean rates of 11.1 subunits s^-1^µM^-1^ ± 0.3 (SE) and 12.7 subunits s^-1^µM^-1^ ± 0.2, respectively (Figure 3C). The mean elongation rate for γ-actin filaments was significantly faster than α-actin, β-actin, or 1:1 mixture of cytoplasmic isoforms (10.9 ± 0.4 subunits s^-1^ µM^-1^; p = 0.02) (Figure 3C).

**Figure 2.**
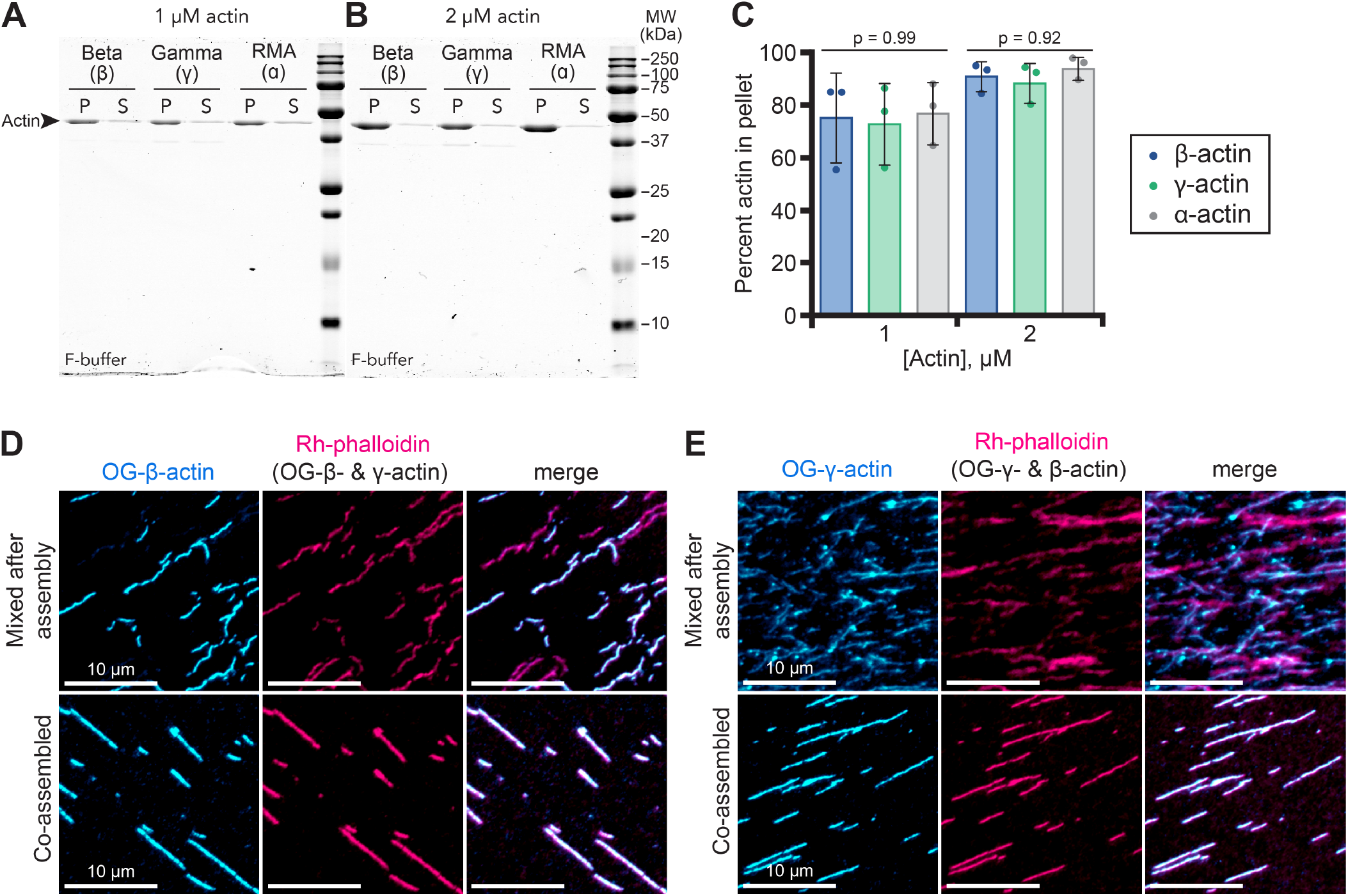
Yeast-made human β- and γ-actin form filaments and co-polymers. **(A-B)** Representative gels of polymerized (P; pellet) or unpolymerized (S; supernatant) actin from pelleting assays performed with (A) 1 or (B) 2 µM actin. Additional uncropped gels shown in Figure S3. **(C)** Quantification by band densitometry of polymerized actin from pellet samples from (A-B)(n = 3 experiments). Statistics: ANOVA; values are not significantly different. **(D)** (top) Mixes of individually polymerized Oregon Green (OG)-labeled β-actin and unlabeled γ-actin stabilized with rhodamine (Rh)-phalloidin or (bottom) co-polymerized OG-β-actin and unlabeled γ-actin, stained with Rh-phalloidin. **(E)** Similar reactions as in (D) with OG-labeled γ-actin and unlabeled β-actin stained with Rh-phalloidin. Scale bars, 10 µm.

**Figure 3.**
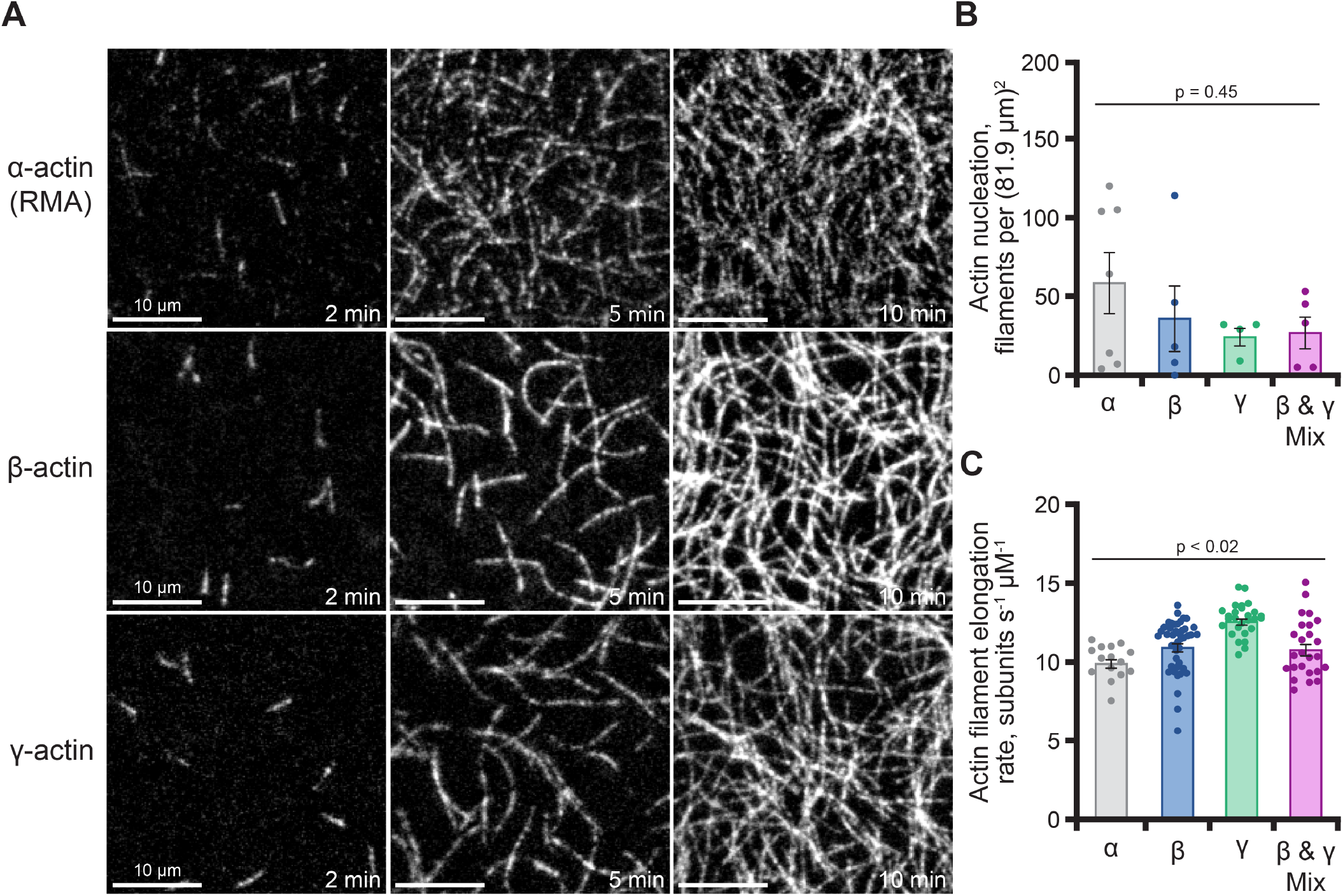
Actin isoforms display similar phases of actin assembly. **(A)** Images from time-lapse TIRF microscopy reactions containing 1 µM total actin monomers as indicated (10% fluorescently labeled RMA). Scale bars, 10 µm. See Movie 1. **(B)** Actin nucleation (n = 3-6) 100 s after the initiation of filament polymerization from reactions as in (A). **(C)** Mean elongation rates for individual filaments (n = 15-45 per condition from at least 3 different reactions; dots) present in reactions as in (A). Statistics: ANOVA; p-values presented are those from ANOVA tests across conditions (all data presented under the line). Only comparisons to γ-actin elongation rates were significantly different (each comparison was p < 0.01 for γ-actin to α, β, or β and γ mix). Comparisons between α- and β-actin were not statistically different in this dataset.

Cellular functions of actin are further modulated through interactions with many regulatory proteins. Therefore, we assessed the activities of several classic regulators of actin polymerization dynamics in the presence of either actin isoform. In mammalian cells, thymosin-β4 (Tβ4) sequesters actin monomers to regulate available subunits for filament polymerization (Skruber et al., 2018; Skruber et al., 2020). We assessed each human actin made in yeast for Tβ4 binding using fluorescence polarization (Figure 4A). GFP-Tβ4 bound β-actin (k_D_ = 1.05 ± 0.30 nM (SE)) and γ-actin (k_D_ = 0.94 ± 0.09 nM) with similar affinities, which were each significantly stronger than skeletal muscle actin (k_D_ = 7.71 nM ± 0.40; p < 0.002) (Figure 4A)(Pimm et al., 2022). This reinforces the notion that actin-binding proteins may have different functions with specific isoforms of actin.

**Figure 4.**
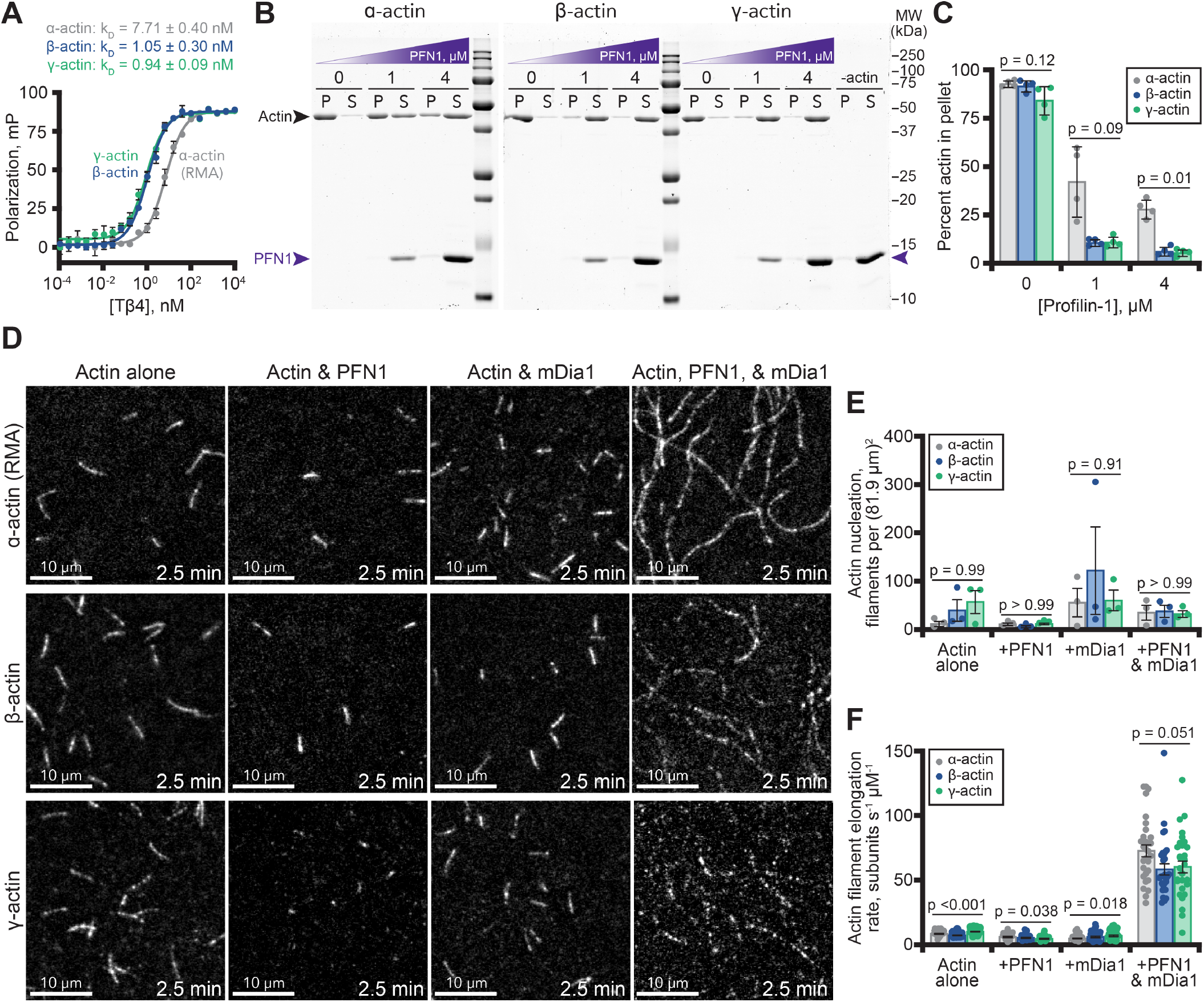
Polymerization of β- or γ-actin with actin assembly factors. **(A)** Fluorescence polarization of GFP-Tβ4 mixed with 10 nM unlabeled α-actin (RMA; grey), β-actin (blue), or γ-actin (green). Curves shown are average fits from n = 3 separate experiments. **(B)** Representative gels displaying the fraction of polymerized (P; pellet) or unpolymerized (S; supernatant) actin present in pelleting assays performed with 2 µM actin and different concentrations of profilin (PFN1; purple arrow)(n = 4). Additional uncropped gels shown in Figure S4. **(C)** Quantification by densitometry of polymerized actin from pellet samples from (B). **(D)** TIRF images from reactions containing 1 µM actin (10% Alexa-488 RMA) and combinations of 2 µM PFN1 and 10 nM mDia1(FH1-C), as indicated. Scale bars, 10 µm. See Movie 2. **(E)** Actin nucleation (n = 3 fields of view; dots) 100 s after the initiation of filament polymerization from reactions as in (D). **(F)** Mean actin elongation rates for individual filaments (n = 30-45 per condition from at least 3 different reactions; dots) present in reactions as in (D). Statistics: ANOVA comparing all conditions; see Figure S5 for re-scaled plot and additional comparisons of the data presented in (F).

To explore this idea further, we assessed the activities of profilin with each isoform of actin. Profilin directly inhibits α-actin assembly (Ferron et al., 2007; Pimm et al., 2022; Skruber et al., 2018). We began using pelleting assays to assess whether profilin preferentially blocked the assembly of β- or γ-actin filaments (Figure 4B). Profilin bound each actin isoform and strongly reduced signals in the pellet fractions of each cytoplasmic actin. We noted that profilin blocks the assembly of β- and γ-actin more effectively than muscle actin (p *≤*0.01)(Figure 4C)(Antoku et al., 2019; Kinosian et al., 2000). However no significant differences were observed for profilin-mediated nucleation in TIRF assays (p > 0.99)(Figure 4D-E).

When present with the actin polymerization stimulating formin (mDia1), profilin enhances actin filament elongation (Chesarone et al., 2010; Li and Higgs, 2005; Valencia and Quinlan, 2021). Thus, to assess whether β- or γ-actin promoted formin-based filament assembly we used TIRF microscopy to monitor actin filament polymerization in the presence of the constitutively active formin, mDia1(FH1-C). Each actin isoform stimulated nucleation to similar levels (p > 0.91)(Figure 4D-E) and stimulated actin filament elongation rates when combined with formin and profilin (Figure 4D, 4F, and S5A-C), with γ-actin filaments elongating significantly faster than the other isoforms (see Figure S5 for all statistical comparisons and rescaled plots). Compared to α-actin filaments which grew at 73.6 ± 4.5 (SE) subunits s_^-1^_ µM_^-1^_, the mean elongation rate for β-actin and γ-actin with profilin and formin was not significantly slower at 59.3 ± 4.2 or 61.2 ± 4.5 subunits s_^-1^_ µM_^-1^_, respectively (p = 0.051)(Figures 4F and S5A-C)(Movie 2). In sum, α-actin from skeletal muscle or either cytoplasmic actin displays similar functions with profilin and the formin mDia1, albeit to different levels.

Finally, the higher-order organization of cellular actin arrays is commonly achieved through the association of proteins that cross-link or bundle filaments. We produced actin bundles with fascin, stained filaments with fluorescent phalloidin, and used TIRF microscopy to compare the higher-order structures made by each isoform (Figure 5A). As actin filaments bundle, the overall area covered by pixel signal decreases (Figure 5A). Meanwhile, the distribution of pixel intensities shifts to brighter pixels (Higaki et al., 2010; Khurana et al., 2010). Thus, we quantitatively measured the extent of actin filament bundling using length and intensity-based metrics to assess total bundling (Figure 5B-C). As actin filaments coalesced into bundles, the number of objects detected in ridge analysis decreased to similar levels, regardless of actin isoform (p = 0.98)(Figure 5B). In contrast, the fluorescence intensity of by β-, γ-, and α-actin filament crosslinked by fascin were similar to each other but each significantly brighter than fascin-absent controls (p < 0.0001)(Figure 5C). In sum, this demonstrates actin filaments of each isoform can be bundled by fascin.

**Figure 5.**
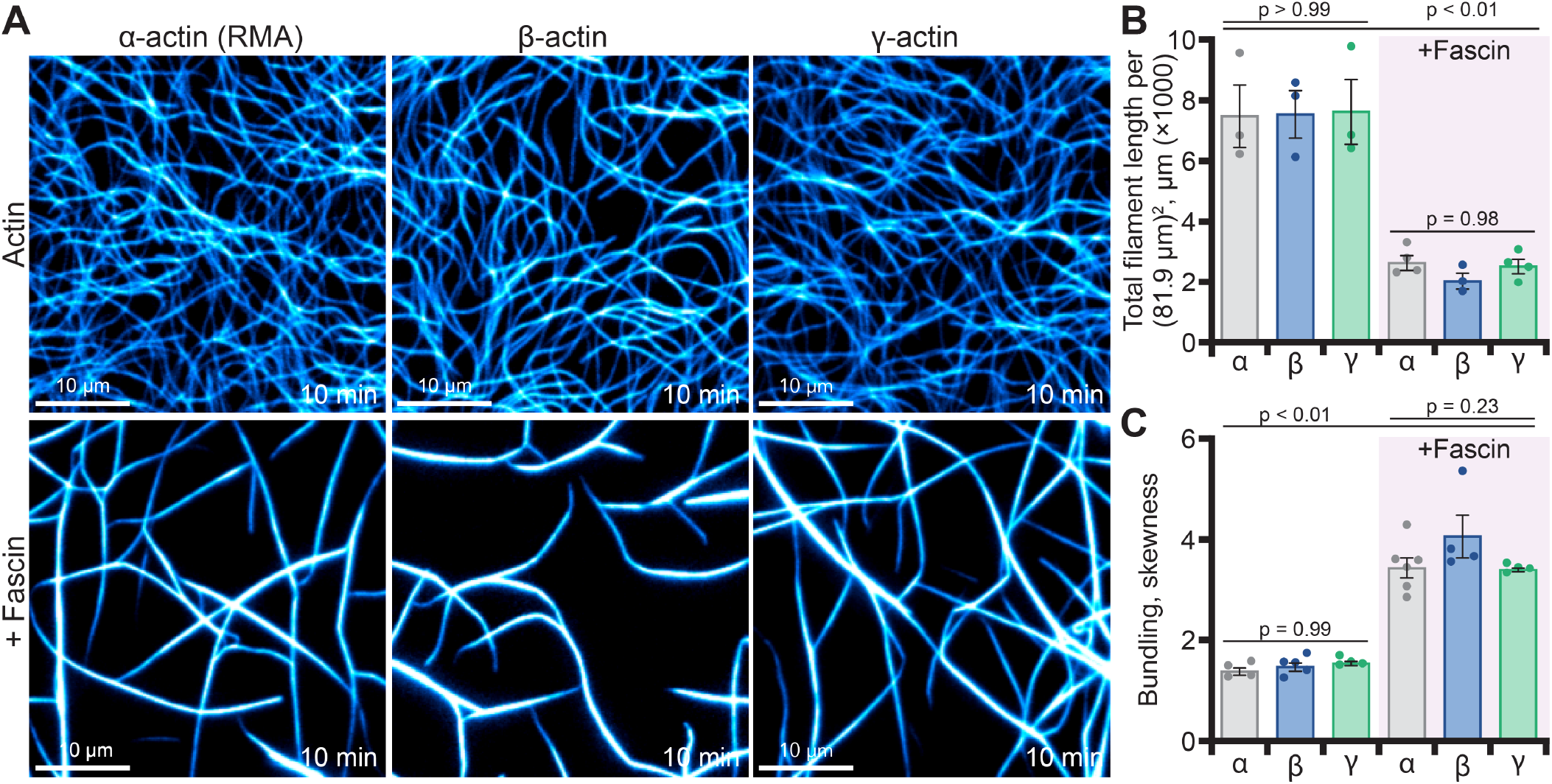
Fascin-mediated actin filament bundling of actin isoforms. **(A)** Representative TIRF images from reactions containing 1 µM unlabeled actin visualized with Alexa-488 phalloidin alone or in the presence of 1 µM fascin. Scale bars, 10 µm. **(B)** Total actin polymer (ridge detection analysis) from reactions as in (A); n = 3-4 fields of view (FOV; dots). **(C)** Filament bundling (skewness) from reactions as in (A); n = 4-6 FOV (dots). Statistics: ANOVA; the only statistical differences found were between untreated controls and fascin-treatments but not between actin isoforms.

## Discussion

We developed budding yeast strains for the expression and purification of human cytoplasmic β- and γ-actin and have demonstrated that these actin isoforms polymerize and interact with several canonical actin binding proteins. There are several benefits to purifying recombinant actin from this system including: yield (0.5-1 mg/L starting culture), no special growth or tissue culture requirements, use of conventional purification reagents and protocols, no additional post-purification processing (e.g., cleavage of fusion tags), and no contaminating “host” actin. Even with these advantages, we note that for both yeast and human actin purifications, a significant amount of protein (>50%) is lost following post-elution dialysis. This likely reflects the denaturing activity of the formamide used to elute actin from the DNaseI column and may be improved by supplementing our approach with an alternative 6×His-gelsolin fragment-based purification (Ceron et al., 2022; Ohki et al., 2009).

Comparisons of actin isoforms or mixes that reflect other species and non-muscle cells are crucial to our understanding of how actin and its regulators truly function. For example, the formin FHOD1 displays differential interactions with actin from different sources, specifically enhancing the nucleation phase of actin assembly with cytoplasmic actin but preventing the assembly of filaments generated from rabbit muscle actin (Antoku et al., 2019; Patel et al., 2018). Profilin and cofilin each bind cytoplasmic actin with higher affinity than muscle actin (Antoku et al., 2019; De La Cruz, 2005; Kinosian et al., 2000). Further, specific myosin motors differentially prefer muscle or non-muscle sources, some even preferring specific β- over γ-actin isoforms for movements (Müller et al., 2013). Our results identify thymosin-β4 as an additional actin regulatory protein influenced by actin source, binding β- or γ-actin with higher affinity than actin purified from muscle.

Over 100 post-translational modifications (PTMs) of cytoplasmic and muscle actins have been described, with common reports of acetylation, arginylation, methylation, and phosphorylation (A et al., 2020; Terman and Kashina, 2013; Varland et al., 2019). Post-translationally modified actin may influence the rate of polymerization, filament-filament interactions, actin-binding protein interactions, cellular localization, and locomotion (Arnesen et al., 2018; Varland et al., 2019; Yamashiro et al., 2014). Perhaps most relevant for the discussion of cytoplasmic actin isoforms is N-terminal acetylation, as β- and γ-actin differ by four residues in this region (Figure 1A). We found that yeast-produced β- and γ-actin were each acetylated on the N-terminal methionine (Table S1, Table S2, and Data File S1)(Cook et al., 1991). One notable missing modification from actin purified in this system is methylation of His73, which is important for the nucleotide-exchange of actin (Wilkinson et al., 2019). Introducing NAA80 or SETD3 to this system may ameliorate these challenges and expand the versatility of our budding yeast-based purification system (Arora et al., 2023; Hatano et al., 2020).

Most mammalian actin is processed to remove the N-terminal methionine and then acetylated or less commonly (and only for β-actin) arginylated at Asp3 (Varland et al., 2019). Studies in yeast and other systems further suggest that N-terminal arginylation occurs on clipped or deacetylated proteins, which may target them for proteasomal degradation (Drazic et al., 2022; Kumar et al., 2016; Nguyen et al., 2019). N-terminal differences in actin isoforms may be important for some applications and less consequential for others (Cook et al., 1992; Hatano et al., 2018). Notably, several studies have successfully purified and utilized versions of human actin lacking N-terminal modifications other than truncations (A et al., 2020; Lu et al., 2015). For those utilizing muscle actin to study interactions with non-muscle actin binding proteins, differences between actin isotypes (primary amino-acid composition) may prove more important than the exact nature of secondary modifications. Our system provides easily obtainable and relatively inexpensive source of bulk cytoplasmic actin that will work well for many applications and that can be useful partners to systems that allow tighter control of specific PTMs or the synthesis of mutated versions of actin (Ceron et al., 2022; Hatano et al., 2020).

## Supplemental information

Supplemental information associated with this work includes: two tables, six figures, two movies, and one dataset.

## Acknowledgements

We thank Stephan Wilkens and Rebecca Oot for the use of their FPLC and assistance with protein labeling. Tom Duncan, Stewart Loh, and Leszek Kotula for the use of equipment. Marcela Alcaide Eligio for assistance with fascin purification. Marc Ridilla for critically reading this work. D.D. Johnson for a greater appreciation of muscle actin. This work was supported by The Research Foundation of SUNY, SUNY Upstate, and NIH grants R01 GM056189 to DCA and R35 GM133485 to JLH-R.

## Author contributions

DCA, BKH, and JLH-R conceived of the project; BKH, JHLR, and MLP performed experiments; EPdJ executed and analyzed mass spectrometry; all authors contributed to writing and editing the manuscript.

## Supplemental Information

### Supplemental Tables

**Table S1.** Mass Spectrometry-based actin secondary modifications.

| Modification<sup>1</sup> | α-actin (RMA)<sup>2</sup> | β-actin | γ-actin |
| --- | --- | --- | --- |
| Acetylation | D1<sup>2</sup>, K50, K68, K215, K238, K284, K291, K315 | M1, K18, K50 | M1, K18, K238 |
| Methylation | K61, K68, H73, K84, H87, H88, H101, N225, N252, K291, N296, N 297, H371 | H40, H87, H88, H101, N128, H161, N252, K291, N296 | N12, K18, H87, H88, H101, H161, N252, K291, N296, H371 |
| Arginylation | H40, D51, W86, F90, A97, K18, T120, T148, L216, C217, V219, A220, C285, K291, D292, L293, E316, Q360 | D3, D4, V10, D51, W86, A97, P98, T120, A135, G150, C217, A220, D292, L299, I329, Q360 | M1, E4, I10, D51, S52, Y69, D80, W86, A97, P98, T120, A135, G150, T160, C217, A220, D292, I329, Q360 |

<sup>1</sup>Data are presented for peptide mass searches for N-terminal or lysine acetylation, methylation, and arginylation.
<sup>2</sup>Numbering of α-actin (RMA) starts at the third amino acid due to the N-terminal truncation of Met-Cys. Thus, Asp3 is listed as D1.

**Table S2.** N-terminal peptide modifications.

| Modification | α-actin (RMA)<sup>1</sup> | β-actin | γ-actin |
| --- | --- | --- | --- |
| Acetylation | D1<sup>1</sup> (4/5) | M1 (11/12)<br>K18 (7/12) | M1 (3/4)<br>K18 (1/4) |
| Arginylation<sup>2</sup> | None detected | D3 (5/9)<br>D4 (3/4) | E4 (1/3) |

<sup>1</sup>Numbering of α-actin (RMA) starts at third amino acid due to N-terminal truncation of Met-Cys.
<sup>2</sup>Arginylation detected as N-terminal modification to truncated peptides.

### Supplemental Figures and Legends

**Figure S1.**
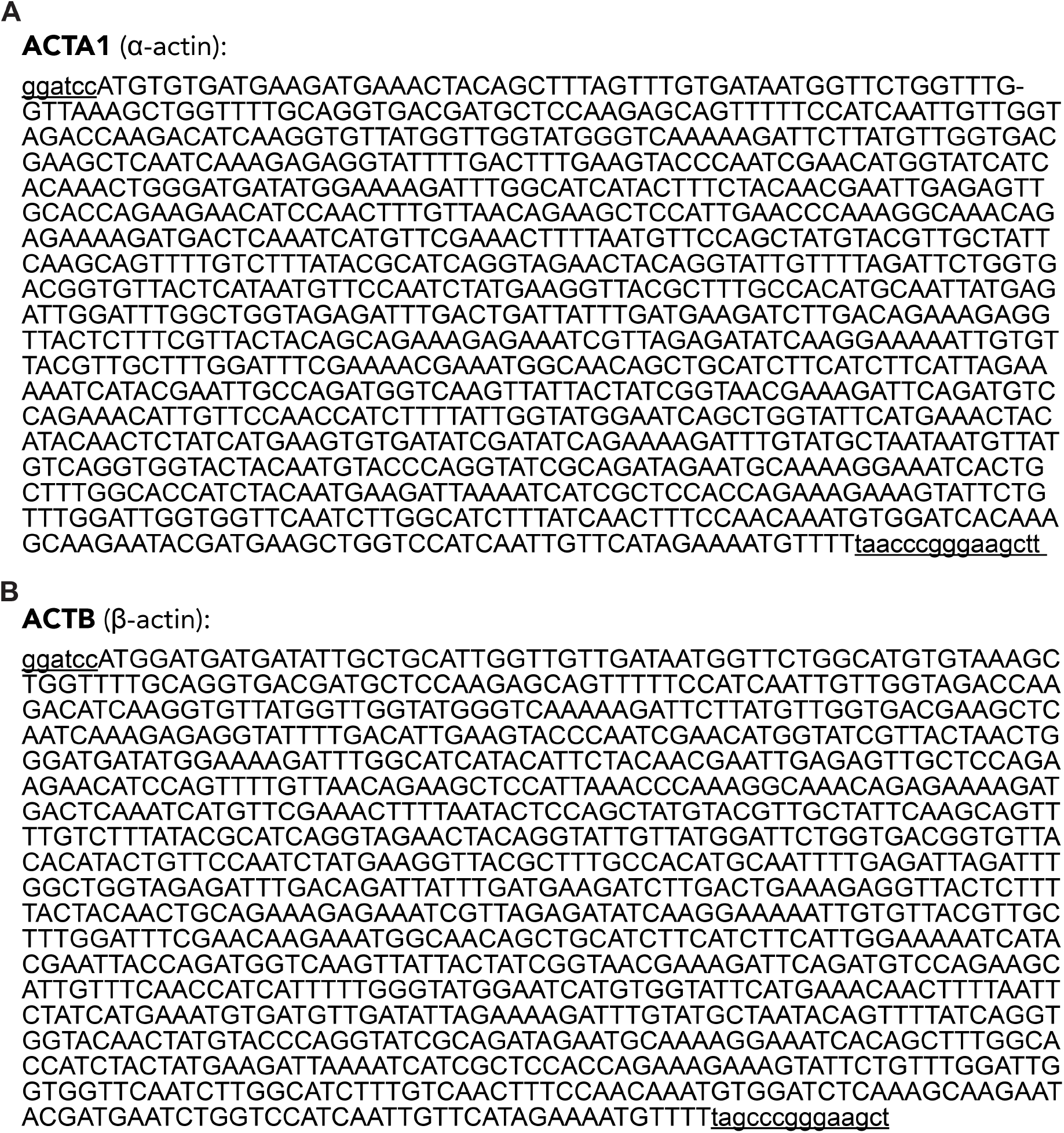
Sequences of yeast-optimized human recombinant actin isoforms. Codon-optimized coding sequences for **(A)** α-actin (ACTA1) and **(B)** β-actin (ACTB) are indicated in upper case letters, whereas flanking sequences including engineered restriction sites are in lower case text.

**Figure S2.**
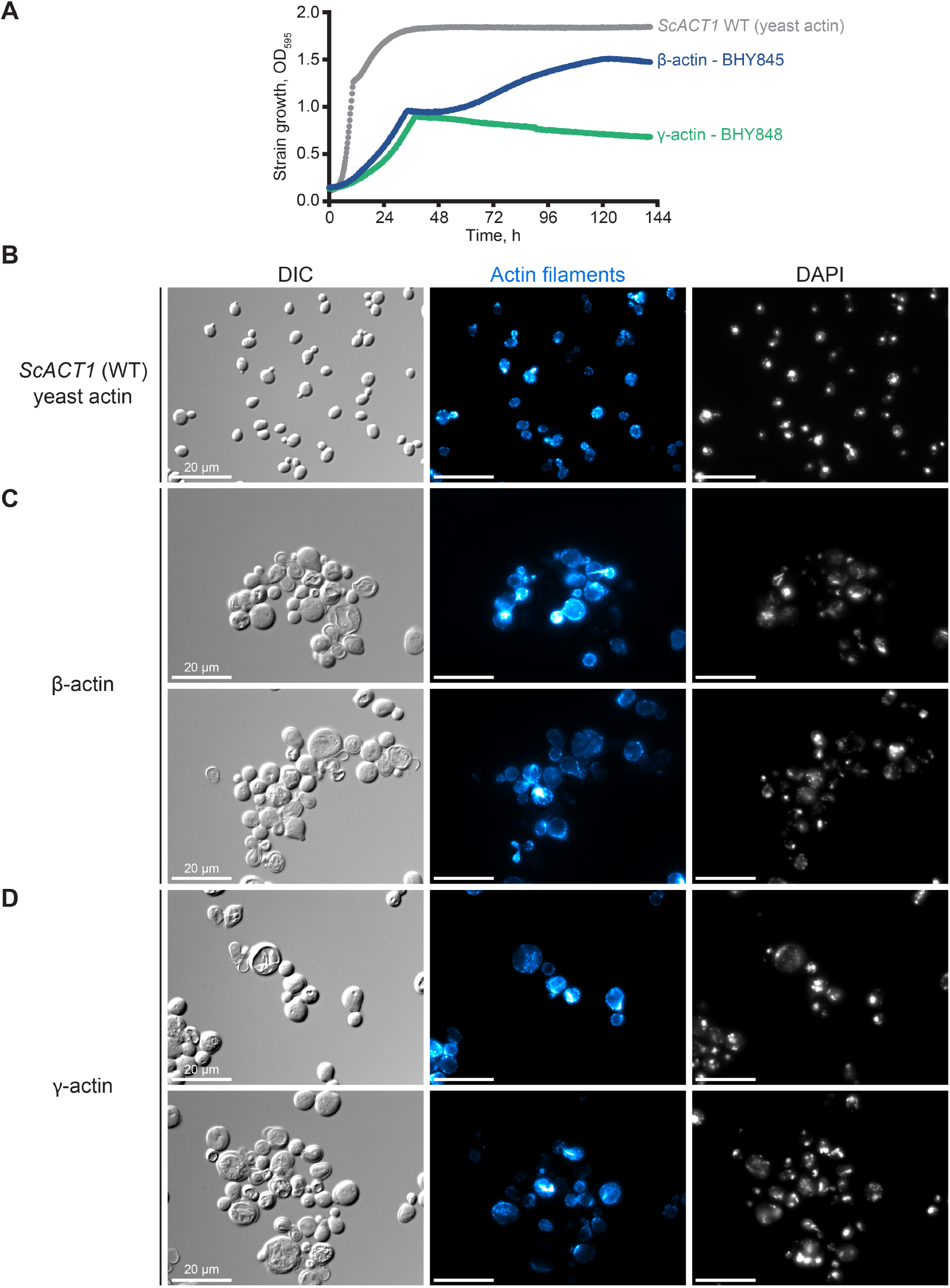
Growth and appearance of yeast expressing α- or γ-actin isoforms. Growth of actin isoform expressing yeast strains performed at 30 °C in YPD media. **(A)** Growth curves of yeast strains expressing yeast actin (*ACT1*; grey), human β-actin (blue), or human γ-actin (green). **(B-D)** Representative DIC and epifluorescence images of yeast expressing (B) yeast actin, (C) human β-actin, or (D) human γ-actin. Left, DIC; middle, rhodamine-phalloidin; right, DAPI. Scale bars, 20 µm.

**Figure S3.**
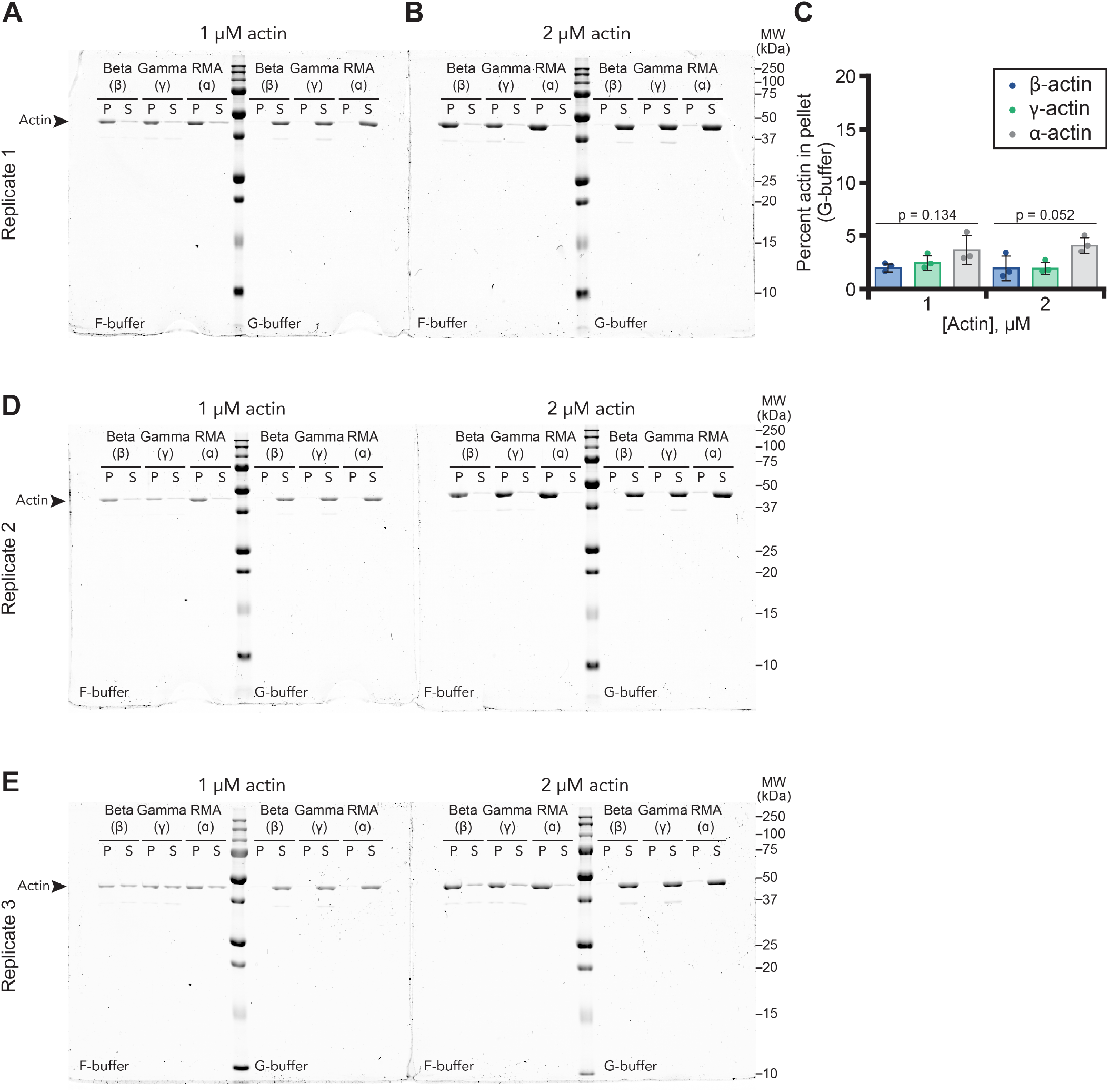
Full gels associated with analysis in Figure 2C. **(A)** Individual gel displaying (P; pellet) or (S; supernatant) fractions from pelleting assays performed with 1 µM actin filaments (F-buffer) or actin monomers (G-buffer). **(B)** Gel as in (A) with 2 µM actin filaments or monomers. Gels shown in A and B are the full gels from Figure 2A and B. **(C)** Quantification by band densitometry of pellet samples from (A-B). Statistics: ANOVA; ns, not significantly different. **(D-E)** Gels from additional replicates for analysis in (C).

**Figure S4.**
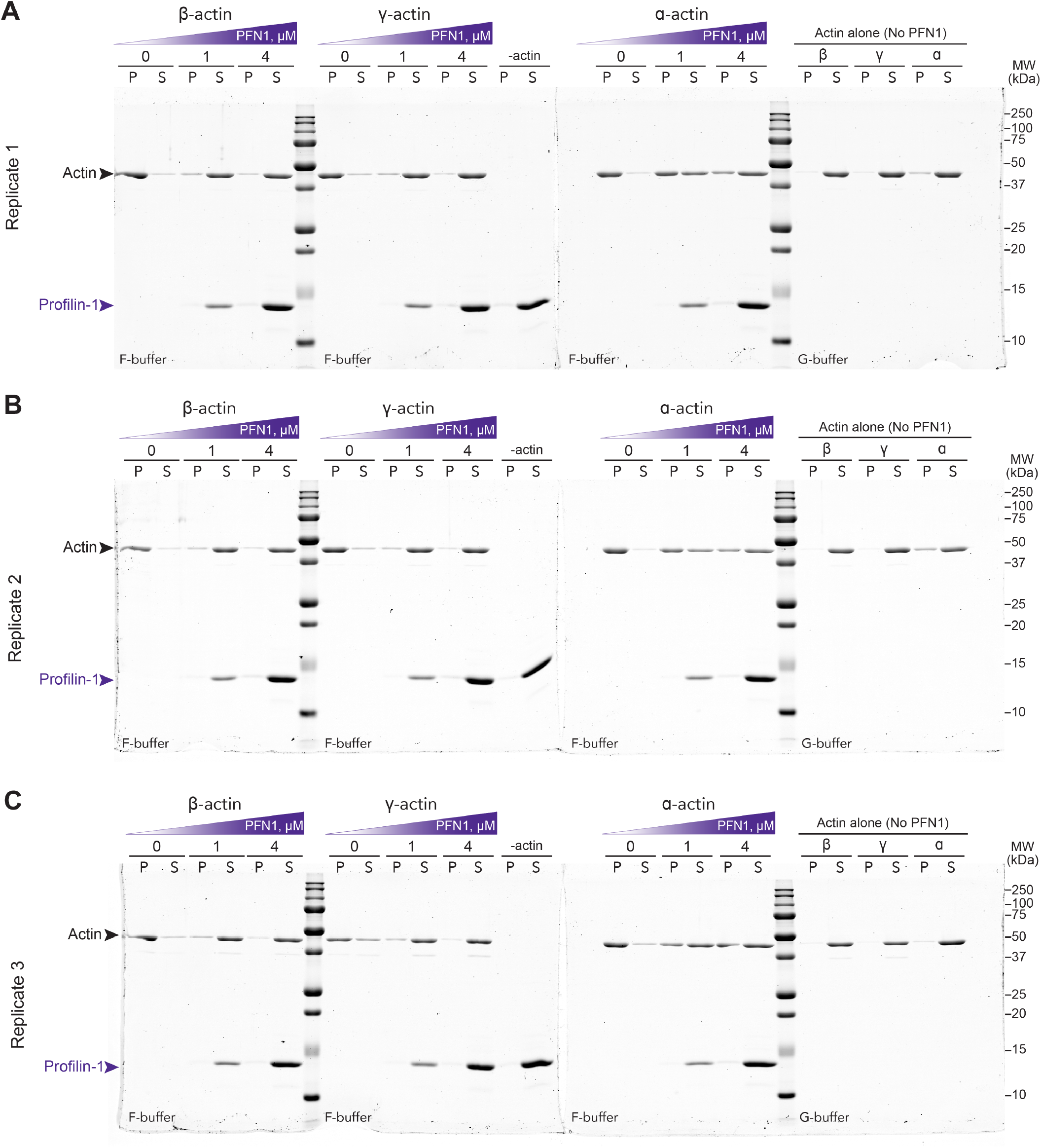
Full gels associated with analysis in Figure 4C. **(A-C)** Full scans of Individual gels displaying (P; pellet) or (S; supernatant) fractions from pelleting assays performed with 2 µM actin filaments (F-actin) and varying amounts of profilin-1 (PFN1; purple arrows). Actin monomer controls (G-buffer) lacking PFN1 are also shown. Panel (A) shows the gels used in Figure 4B.

**Figure S5.**
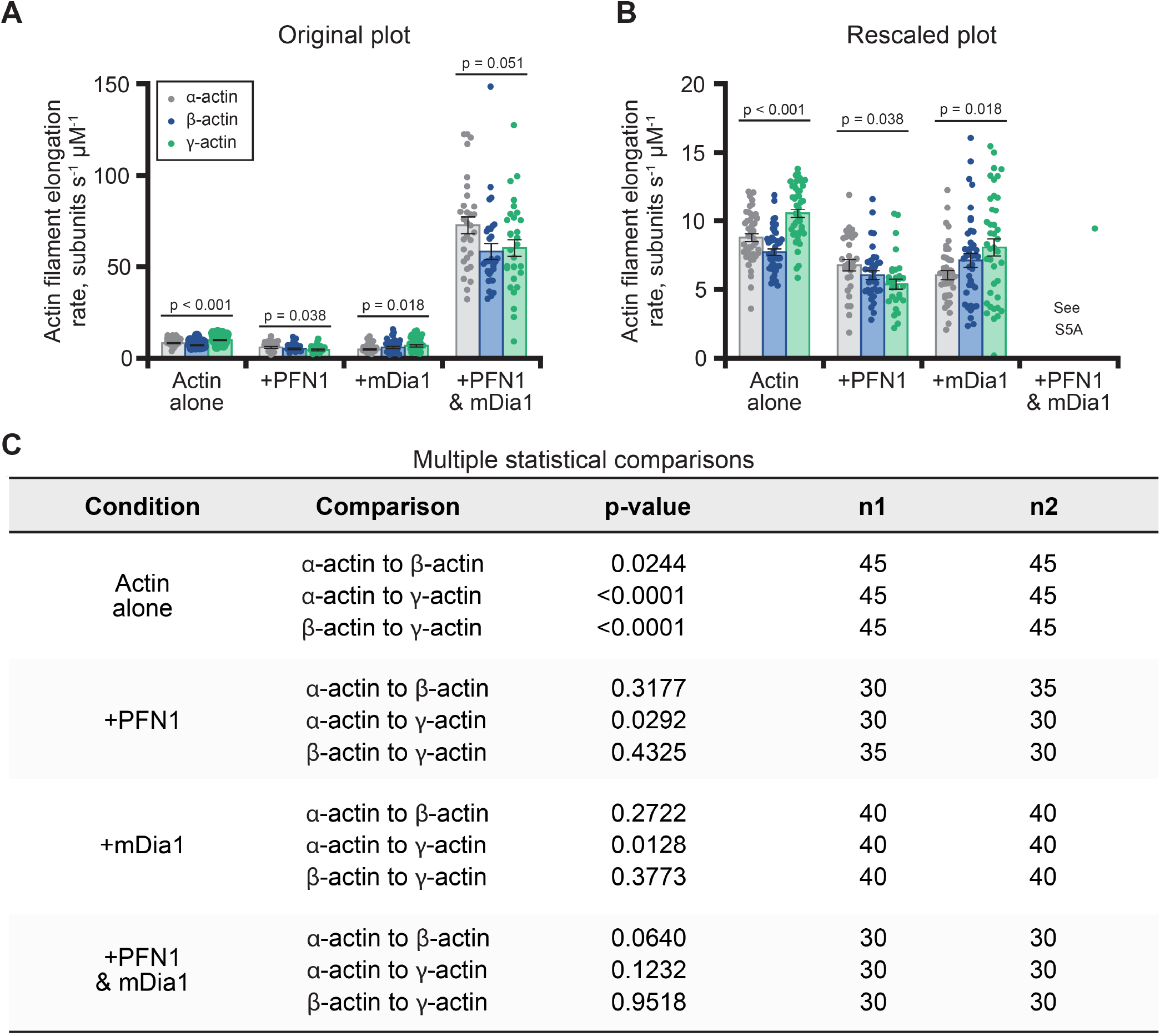
Comparison of Figure 4F and re-scaled version to view slower elongation rates. **(A)** Original plot of actin filament elongation rates from Figure 4F. **(B)** Data from Figure 4F or panel (A) rescaled to make the distributions of slower elongation rates more visible. Statistics: ANOVA comparison of all treatments under the line. **(C)** Table displaying additional individual statistical comparisons.

**Figure S6.**
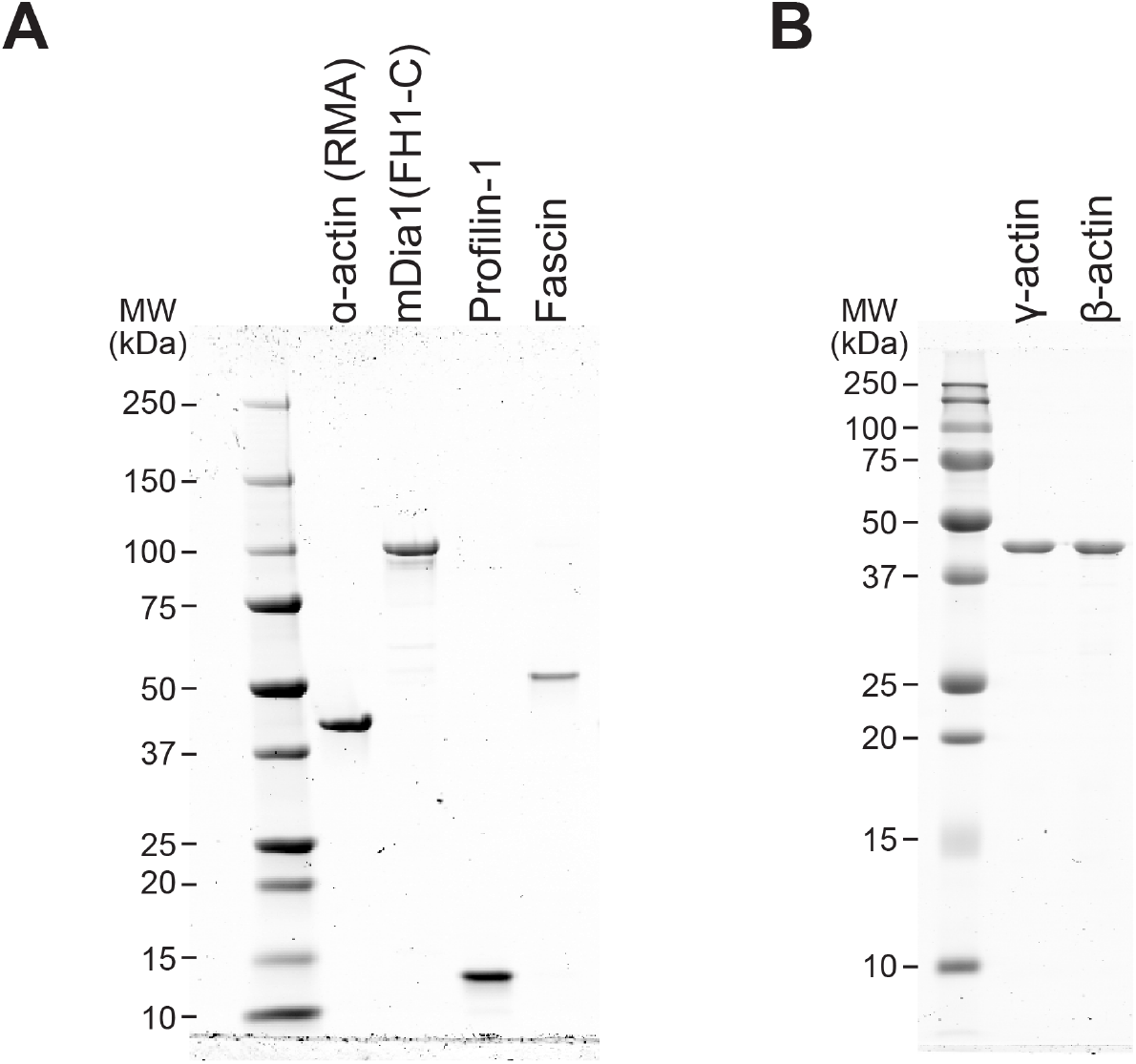
Purity of proteins used in TIRF assays. Coomassie-stained gels showing the purity of proteins used in this study, as follows: **(A)** 360 ng α-actin (RMA), 375 ng mDia1(FH1-C), 165 ng profilin-1, 280 ng fascin, **(B)** 390 ng β-actin, and 310 ng γ-actin.

### Movie Legends

**Movie 1**. β- **and** γ**-actin form actin filament polymers**. TIRF movies comparing the polymerization of actin isoforms from reactions containing: 1 µM actin of each isoform (10% Alexa-488 RMA). Scale bar, 20 µm. Playback, 10 fps.

**Movie 2. The formin mDia1 can nucleate and rapidly elongate** β**- and** γ **-actin filaments**. TIRF movies of reactions containing labeled combinations of 1 µM α-actin, β-actin, or γ-actin (10% Alexa-488 RMA),10 nM mDia1(FH1-C), and 2 µM profilin-1. Scale bar, 20 µm. Playback, 10 fps.

## References

1. A M., Fung T. S., Francomacaro, L. M., Huynh, T., Kotila, T., Svindrych, Z. and Higgs, H. N. (2020). Regulation of INF2-mediated actin polymerization through site-specific lysine acetylation of actin itself. Proceedings of the National Academy of Sciences. U.S.A. 117: 439–447.

2. Aggeli, D., E. Kish-Trier, M.C. Lin, B. Haarer, G. Cingolani, J.A. Cooper, S. Wilkens, and D.C. Amberg. 2014. Coordination of the filament stabilizing versus destabilizing activities of cofilin through its secondary binding site on actin. Cytoskeleton. 71: 361–379.

3. Allen, P.G., C.B. Shuster, J. Kas, C. Chaponnier, P.A. Janmey, and I.M. Herman. 1996. Phalloidin binding and rheological differences among actin isoforms. Biochemistry. 35: 14062–14069.

4. Antoku, S., W. Wu, L.C. Joseph, J.P. Morrow, H.J. Worman, and G.G. Gundersen. 2019. ERK1/2 Phosphorylation of FHOD Connects signaling and nuclear positioning alternations in cardiac laminopathy. Developmental Cell. 51: 602–616e612.

5. Arnesen, T., R. Marmorstein, and R. Dominguez. 2018. Actin ‘s N-terminal acetyltransferase uncovered. Cytoskeleton. 75: 318–322.

6. Arora, A. S., Huang, H.-L., Singh, R., Narui, Y., Suchenko, A., Hatano, T., Heissler, S. M., Balasubramanian, M. K. and Chinthalapudi, K. (2023). Structural insights into actin isoforms. eLife. 12: e82015.

7. Bergeron, S.E., M. Zhu, S.M. Thiem, K.H. Friderici, and P.A. Rubenstein. 2010. Ion-dependent polymerization differences between mammalian beta- and gamma-nonmuscle actin isoforms. Journal of Biological Chemistry. 285: 16087–16095.

8. Bookwalter, C.S., and K.M. Trybus. 2006. Functional consequences of a mutation in an expressed human alpha-cardiac actin at a site implicated in familial hypertrophic cardiomyopathy. Journal of Biological Chemistry. 281: 16777–16784.

9. Brademan, D. R., Riley, N. M., Kwiecien, N. W. and Coon, J. J. (2019). Interactive peptide spectral annotator: a versatile web-based tool for proteomic applications. Molecular Cell Proteomics. 18: S193–S201.

10. Ceron, R. H., Carman, P. J., Rebowski, G., Boczkowska, M., Heuckeroth, R. O. and Dominguez, R. (2022). A solution to the long-standing problem of actin expression and purification. Proceedings of the National Academy of Sciences. U.S.A. 119: e2209150119.

11. Chesarone, M.A., A.G. DuPage, and B.L. Goode. 2010. Unleashing formins to remodel the actin and microtubule cytoskeletons. Nature Reviews Molecular Cell Biology. 11: 62–74.

12. Chin, S. M., Hatano, T., Sivashanmugam, L., Suchenko, A., Kashina, A. S., Balasubramanian, M. K. and Jansen, S. (2022). N-terminal acetylation and arginylation of actin determines the architecture and assembly rate of linear and branched actin networks. Journal of Biological Chemistry. 298: 102518.

13. Cook, R.K., W.T. Blake, and P.A. Rubenstein. 1992. Removal of the amino-terminal acidic residues of yeast actin. Studies in vitro and in vivo. Journal of Biological Chemistry. 267: 9430–9436.

14. Cook, R.K., D.R. Sheff, and P.A. Rubenstein. 1991. Unusual metabolism of the yeast actin amino terminus. Journal of Biological Chemistry. 266: 16825–16833.

15. De La Cruz, E.M. 2005. Cofilin binding to muscle and non-muscle actin filaments: isoform-dependent cooperative interactions. Journal of Molecular Biology. 346: 557–564.

16. Drazic, A., Timmerman, E., Kajan, U., Marie, M., Varland, S., Impens, F., Gevaert, K. and Arnesen, T. (2022). The final maturation state of β-actin involves N-terminal acetylation by NAA80, not N-terminal arginylation by ATE1. Journal of Molecular Biology. 434: 167397.

17. Ferron, F., G. Rebowski, S.H. Lee, and R. Dominguez. 2007. Structural basis for the recruitment of profilin-actin complexes during filament elongation by Ena/VASP. EMBO Journal. 26: 4597–4606.

18. Geissler, S., K. Siegers, and E. Schiebel. 1998. A novel protein complex promoting formation of functional alpha- and gamma-tubulin. EMBO Journal. 17: 952–966.

19. Grantham, J. 2020. The molecular chaperone CCT/TRiC: an essential component of proteostasis and a potential modulator of protein aggregation. Frontiers Genetics. 11: 172.

20. Hatano, T., S. Alioto, E. Roscioli, S. Palani, S.T. Clarke, A. Kamnev, J.R. Hernandez-Fernaud, L. Sivashanmugam, Y.L.B. Chapa, A.M.E. Jones, R.C. Robinson, K. Sampath, M. Mishima, A.D. McAinsh, B.L. Goode, and M.K. Balasubramanian. 2018. Rapid production of pure recombinant actin isoforms in Pichia pastoris. Journal of Cell Science. 131: jcs213827.

21. Hatano, T., L. Sivashanmugam, A. Suchenko, H. Hussain, and M.K. Balasubramanian. 2020. Pick-ya actin -a method to purify actin isoforms with bespoke key post-translational modifications. Journal of Cell Science. 133: jcs241406.

22. Henty-Ridilla, J.L. 2022. Visualizing actin and microtubule coupling dynamics in vitro by total internal reflection fluorescence (TIRF) microscopy. Journal of Visual Experiments. 185: e64074.

23. Hertzog, M. and Carlier, M. (2005). Functional characterization of proteins regulating actin assembly. Current Protocols in Cell Biology. 26: 13.6.1-13.6.23.

24. Higaki, T., N. Kutsuna, T. Sano, N. Kondo, and S. Hasezawa. 2010. Quantification and cluster analysis of actin cytoskeletal structures in plant cells: role of actin bundling in stomatal movement during diurnal cycles in Arabidopsis guard cells. Plant Journal. 61: 156–165.

25. Hundt, N., M. Preller, O. Swolski, A.M. Ang, H.G. Mannherz, D.J. Manstein, and M. Muller. 2014. Molecular mechanisms of disease-related human betaactin mutations p.R183W and p.E364K. FEBS Journal. 281: 5279–5291.

26. Jansen, S., A. Collins, C. Yang, G. Rebowski, T. Svitkina, and R. Dominguez. 2011. Mechanism of actin filament bundling by fascin. Journal of Biological Chemistry. 286: 30087–30096.

27. Kalhor, H. R., Niewmierzycka, A., Faull, K. F., Yao, X., Grade, S., Clarke, S. and Rubenstein, P. A. (1999). A Highly Conserved 3-Methylhistidine Modification is absent in yeast actin. Archives of Biochemistry and Biophysics. 370: 105–111.

28. Karakozova, M., M. Kozak, C.C. Wong, A.O. Bailey, J.R. Yates, 3rd, A. Mogilner, H. Zebroski, and A. Kashina. 2006. Arginylation of beta-actin regulates actin cytoskeleton and cell motility. Science. 313: 192–196.

29. Karlsson, R. 1988. Expression of chicken beta-actin in Saccharomyces cerevisiae. Gene. 68: 249–257.

30. Karlsson, R., P. Aspenstrom, and A.S. Bystrom. 1991. A chicken beta-actin gene can complement a disruption of the Saccharomyces cerevisiae ACT1 gene. Molecular and Cellular Biology. 11: 213–217.

31. Khurana, P., J.L. Henty, S. Huang, A.M. Staiger, L. Blanchoin, and C.J. Staiger. 2010. Arabidopsis VILLIN1 and VILLIN3 have overlapping and distinct activities in actin bundle formation and turnover. Plant Cell. 22: 2727–2748.

32. Kijima, S.T., K. Hirose, S.G. Kong, M. Wada, and T.Q. Uyeda. 2016. Distinct biochemical properties of Arabidopsis thaliana actin isoforms. Plant Cell Physiology. 57: 46–56.

33. Kinosian, H.J., L.A. Selden, L.C. Gershman, and J.E. Estes. 2000. Interdependence of profilin, cation, and nucleotide binding to vertebrate non-muscle actin. Biochemistry. 39: 13176–13188.

34. Kuhn, J. R. and Pollard, T. D. (2005). Real-time measurements of actin filament polymerization by total internal reflection fluorescence microscopy. Biophysical Journal. 88: 1387–1402.

35. Kumar, A., Birnbaum, M. D., Patel, D. M., Morgan, W. M., Singh, J., Barrientos, A. and Zhang, F. (2016). Posttranslational arginylation enzyme Ate1 affects DNA mutagenesis by regulating stress response. Cell Death and Disease. 7: e2378–e2378.

36. Li, F., and H.N. Higgs. 2003. The mouse Formin mDia1 is a potent actin nucleation factor regulated by autoinhibition. Current Biology. 13: 1335–1340.

37. Li, M.-M., Nilsen, A., Shi, Y., Fusser, M., Ding, Y.-H., Fu, Y., Liu, B., Niu, Y., Wu, Y.-S., Huang, C.-M., et al. (2013). ALKBH4-dependent demethylation of actin regulates actomyosin dynamics. Nature Communications. 4: 1832.

38. Liu, X., M.L. Pimm, B. Haarer, A.T. Brawner, and J.L. Henty-Ridilla. 2022. Biochemical characterization of actin assembly mechanisms with ALS-associated profilin variants. European Journal of Cell Biology. 101: 151212.

39. Lu, H., Fagnant, P. M., Bookwalter, C. S., Joel, P. and Trybus, K. M. (2015). Vascular disease-causing mutation R258C in ACTA2 disrupts actin dynamics and interaction with myosin. Proceedings of the National Academy of Sciences. U.S.A. 112: E4168–E4177.

40. McKane, M., Wen, K.-K., Boldogh, I. R., Ramcharan, S., Pon, L. A. and Rubenstein, P. A. (2005). A mammalian actin substitution in yeast actin (H372R) causes a suppressible mitochondria/vacuole phenotype. Journal Biological Chemistry. 280: 36494–36501.

41. McKane, M., Wen, K.-K., Meyer, A. and Rubenstein, P. A. (2006). Effect of the substitution of muscle actin-specific subdomain 1 and 2 residues in yeast actin on actin function. Journal Biological Chemistry. 281: 29916–29928.

42. Millan-Zambrano, G., and S. Chavez. 2014. Nuclear functions of prefoldin. Open Biology. 4.

43. Moradi, M., R. Sivadasan, L. Saal, P. Luningschror, B. Dombert, R.J. Rathod, D.C. Dieterich, R. Blum, and M. Sendtner. 2017. Differential roles of alpha-, beta-, and gamma-actin in axon growth and collateral branch formation in motoneurons. Journal of Cell Biology. 216: 793–814.

44. Muller, M., R.P. Diensthuber, I. Chizhov, P. Claus, S.M. Heissler, M. Preller, M.H. Taft, and D.J. Manstein. 2013. Distinct functional interactions between actin isoforms and nonsarcomeric myosins. PLoS ONE. 8: e70636.

45. Muller, M., A.J. Mazur, E. Behrmann, R.P. Diensthuber, M.B. Radke, Z. Qu, C. Littwitz, S. Raunser, C.A. Schoenenberger, D.J. Manstein, and H.G. Mannherz. 2012. Functional characterization of the human alpha-cardiac actin mutations Y166C and M305L involved in hypertrophic cardiomyopathy. Cellular and Molecular Life Sciences. 69: 3457–3479.

46. Namba, Y., M. Ito, Y. Zu, K. Shigesada, and K. Maruyama. 1992. Human T cell L-plastin bundles actin filaments in a calcium-dependent manner. Journal of Biochemistry. 112: 503–507.

47. Nguyen, K. T., Kim, J.-M., Park, S.-E. and Hwang, C.-S. (2019). N-terminal methionine excision of proteins creates tertiary destabilizing N-degrons of the Arg/Nend rule pathway. Journal of Biological Chemistry. 294: 4464–4476.

48. Noguchi, T.Q., N. Kanzaki, H. Ueno, K. Hirose, and T.Q. Uyeda. 2007. A novel system for expressing toxic actin mutants in Dictyostelium and purification and characterization of a dominant lethal yeast actin mutant. Journal of Biological Chemistry. 282: 27721–27727.

49. Ohki, T., C. Ohno, K. Oyama, S.V. Mikhailenko, and S. Ishiwata. 2009. Purification of cytoplasmic actin by affinity chromatography using the C-terminal half of gelsolin. Biochemical and Biophysical Research Communications. 383: 146–150.

50. Parker, F., Baboolal, T. G. and Peckham, M. (2020). Actin mutations and their role in disease. International Journal of Molecular Sciences. 21: 3371.

51. Patel, A.A. Z.A. Oztug Durer, A.P. van Loon, K.V. Bremer, and M.E. Quinlan. 2018. Drosophila and human FHOD family formin proteins nucleate actin filaments. Journal of Biological Chemistry. 293:532-540.

52. Perrin, B.J., and J.M. Ervasti. 2010. The actin gene family: function follows isoform. Cytoskeleton. 67: 630–634.

53. Pimm, M.L., J. Hotaling, and J.L. Henty-Ridilla. 2020. Profilin choreographs actin and microtubules in cells and cancer. International Review of Cell and Molecular Biology. 355:155–204.

54. Rappsilber, J., Y. Ishihama, and M. Mann. 2003. Stop and go extraction tips for matrix-assisted laser desorption/ionization, nanoelectrospray, and LC/MS sample pretreatment in proteomics. Analytical Chemistry. 75: 663–670.

55. Rutkevich, L.A., D.J. Teal, and J.F. Dawson. 2006. Expression of actin mutants to study their roles in cardiomyopathy. Canadian Journal Physiology Pharmacology. 84: 111–119.

56. Schafer, D. A., Jennings, P. B. and Cooper, J. A. (1998). Rapid and efficient purification of actin from nonmuscle sources. Cell Motility and Cytoskeleton. 39: 166–171.

57. Schindelin, J., I. Arganda-Carreras, E. Frise, V. Kaynig, M. Longair, T. Pietzsch, S. Preibisch, C. Rueden, S. Saalfeld, B. Schmid, J.Y. Tinevez, D.J. White, V. Hartenstein, K. Eliceiri, P. Tomancak, and A. Cardona. 2012. Fiji: an open-source platform for biological-image analysis. Nature Methods. 9: 676–682.

58. Skruber, K., P.V. Warp, R. Shklyarov, J.D. Thomas, M.S. Swanson, J.L. Henty-Ridilla, T.A. Read, and E.A. Vitriol. 2020. Arp2/3 and Mena/VASP require Profilin 1 for actin network assembly at the leading edge. Current Biology. 30:2651–2664 e2655.

59. Steger, C. 1998. An unbiased detector of curvilinear structures. IEEE Transactions on Pattern Analysis and Machine Intelligence. 20: 113–125.

60. Tamura, M., K. Ito, S. Kunihiro, C. Yamasaki, and M. Haragauchi. 2011. Production of human beta-actin and a mutant using a bacterial expression system with a cold shock vector. Protein Expression and Purification. 78: 1–5.

61. Terman, J.R., and A. Kashina. 2013. Post-translational modification and regulation of actin. Current Opinion in Cell Biology. 25: 30–38.

62. Valencia, D.A., and M.E. Quinlan. 2021. Formins. Current Biology. 31: R517–R522.

63. Valpuesta, J.M., J. Martin-Benito, P. Gomez-Puertas, J.L. Carrascosa, and K.R. Willison. 2002. Structure and function of a protein folding machine: the eukaryotic cytosolic chaperonin CCT. FEBS letters. 529: 11–16.

64. Varland, S., J. Vandekerckhove, and A. Drazic. 2019. Actin post-translational modifications: the cinderella of cytoskeletal control. Trends in Biochemical Sciences. 44: 502–516.

65. von der Ecken, J., S.M. Heissler, S. Pathan-Chhatbar, D.J. Manstein, and S. Raunser. 2016. Cryo-EM structure of a human cytoplasmic actomyosin complex at near-atomic resolution. Nature. 534: 724–728.

66. Wagner, T., M. Hiner, and Xraynaud. 2017. Thorstenwagner/Ij-Ridgedetection: ridge detection 1.4.0. Zenodo.

67. Wilkinson, A. W., Diep, J., Dai, S., Liu, S., Ooi, Y. S., Song, D., Li, T.-M., Horton, J. R., Zhang, X., Liu, C., et al. (2019). SETD3 is an actin histidine methyltransferase that prevents primary dystocia. Nature. 565: 372–376.

68. Xu, T., Wong, C. C. L., Kashina, A. and Yates, J. R. (2009). Identification of N-terminally arginylated proteins and peptides by mass spectrometry. Nature Protocols. 4: 325–332.

69. Yamashiro, S., D.S. Gokhin, Z. Sui, S.E. Bergeron, P.A. Rubenstein, and V.M. Fowler. 2014. Differential actin-regulatory activities of Tropomodulin1 and Tropomodulin3 with diverse tropomyosin and actin isoforms. Journal of Biological Chemistry. 289: 11616–11629.

